## Supplemental Figure 1a for "Functional gene categories differentiate maize leaf drought-related microbial epiphytic communities"

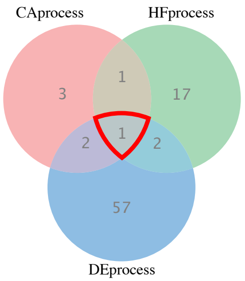


nGO:0006351 transcription, DNA-templated

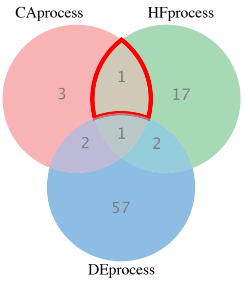


GO:0006865 amino acid transport

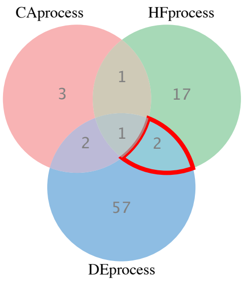


GO:0006355 regulation of transcription, DNA-templated
GO:0006412 translation

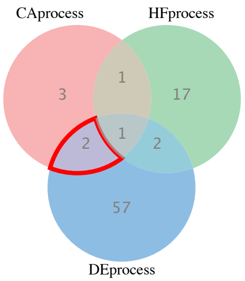


GO:0006457 protein folding
GO:0009097 isoleucine biosynthetic process
