## Supplemental Figure 1b for "Functional gene categories differentiate maize leaf drought-related microbial epiphytic communities"

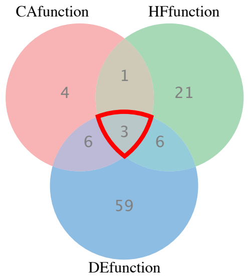


GO:0003677 DNA binding
GO:0022857 transmembrane transporter activity
GO:0046872 metal ion binding

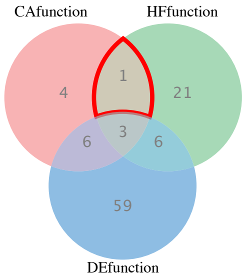


GO:0005525 GTP binding

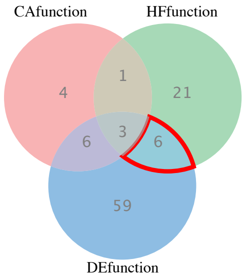


GO:0000287 magnesium ion binding
GO:0004519 endonuclease activity
GO:0005524 ATP binding
GO:0008270 zinc ion binding
GO:0009055 electron carrier activity
GO:00515394 iron, 4 sulfur cluster binding

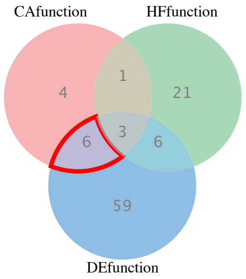


GO:0003700 sequence-specific DNA binding transcription factor activity
GO:0003755 peptidyl-prolyl cis-trans isomerase activity
GO:0004455 ketol-acid reductoisomerase activity
GO:0016787 hydrolase activity
GO:0050661 NADP binding
GO:0051287 NAD binding
