## Supplementary figures and images for "Functional gene categories differentiate maize leaf drought-related microbial epiphytic communities"

### Supplemental Figure2

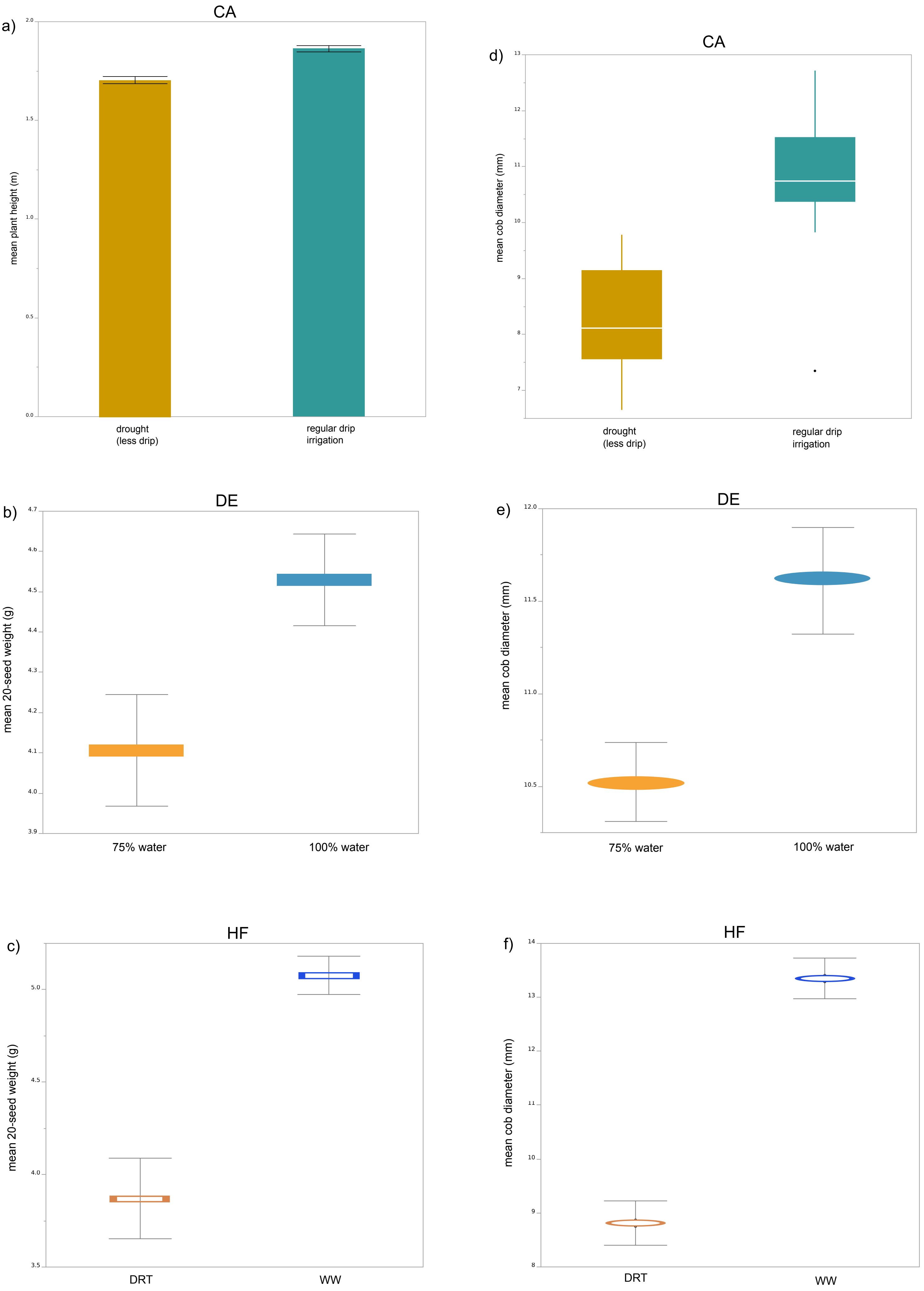
