## Supplemental File S2_7 for "Functional gene categories differentiate maize leaf drought-related microbial epiphytic communities"

Supplemental File 7 plant trait data readme and metadata

Abbreviations:

CA is the California-Albany field site, 37°53'12.8"N 122°17'59.8"W

DE is the Texas Dumas Etter field site, 35.998744°N 101.988583°W

HF is the Texas Halfway field site, 34.184136°N 101.943636°W

Filenames and metadata on file contents:

Supplemental File 1 CA cob_diameter.xls

Column 1 header Treatment, can have values of: drought (less drip) or regular drip irrigation

Column 2 header cob_diameter_mm, contains cob diameters in millimeters

148 rows

Supplemental File 2 CA plant_height.xls

Column 1 header treatment	has values of: dry or wet

Column 2 header entry is a plot number specific to that field; plant height measurements were taken in all plants in each entry plot

Column 2 header plant_height_in is plant height recorded in inches

80 rows

Supplemental File 3 DE cob_diameter.xls

Column 1 header treatment	can have the value wet or dry

Column 2 header entry	is a plot number specific to that field

Column 3 header cob_diameter_mm, contains cob diameters in millimeters

37 rows

Supplemental File 4 DE seed wt.xls

Column 1 header treatment, can have values d (for dry) or w (for wet)

Column 2 header 20_seed_weight_g, weight of packet with 20 seeds, in grams

57 rows

Supplemental File 5 HF cob_diameter.xls

Column 1 header treatment, can have values WW (for wet) and DRT (for dry)

Column 2 header entry, this is a plot number specific to that field

Column 3 header cob_diameter_mm, contains cob diameters measurements in millimeters

36 rows

Supplemental File 6 HF seed wt.xls

Column 1 header treatment with valued d (for dry) or w (for wet)

Column 2 header 20_seed_weight_g, weight of packet with 20 seeds, in grams

33 rows
